# Artificial diets modulate infection rates by *Nosema ceranae* in bumblebees

**DOI:** 10.1101/2020.02.27.967455

**Authors:** Tamara Gómez-Moracho, Tristan Durand, Cristian Pasquaretta, Philipp Heeb, Mathieu Lihoreau

**Author notes:** Author for correspondence: Mathieu Lihoreau.

## Abstract

Parasites alter the physiology and behaviour of their hosts. In domestic honey bees, the microsporidia *Nosema ceranae* induces an energetic stress and impairs the behaviour of foragers, potentially leading to colony collapse. Whether this emerging parasite similarly affects wild pollinators is little understood because of the low success rates of experimental infection protocols. Here we established a new apporach for infecting bumblebees (*Bombus terrestris*) with controlled amounts of *N. ceranae*, by briefly exposing individual bumblebees to a sucrose solution containing parasite spores, before feeding them with artificial diets. We validated our protocol by testing the effect of two spore dosages and two diets varying in their protein to carbohydrate ratio, on the prevalence of the parasite (proportion of infected bees), the intensity of infection (spore count in the gut), and the survival of bumblebees. Insects fed a low-protein high-carbohydrate diet showed highest parasite prevalence (up to 70%) but lived longest, suggesting that immunity and survival of bumblebees are maximised at different protein to carbohydrate ratios. Spore dosage had no effect on parasite infection rate and host survival. The identification of experimental conditions for successfully infecting bumblebees with *N. ceranae* in the lab will facilitate investigations of the sub-lethal effects of this parasite on the behaviour and cognition of wild pollinators.

## 1. Introduction

Bees face a high diversity of parasites and pathogens that negatively affect their metabolism (Li et al., 2018; Mayack and Naug, 2009), immune system (Antúnez et al., 2009; Li et al., 2017), behaviour and cognition (Gómez-Moracho et al., 2017), altogether compromising the fitness of individuals and colonies (Goulson et al. 2015; Klein et al., 2017).

The microsporidia *Nosema ceranae* is one of the most prevalent parasites of honey bees worldwide and is thought to be a major cause of colony declines (Martín-Hernández et al., 2018). *N. ceranae* invades the epithelial cells of the honey bee midgut where it replicates (García-Palencia et al., 2010; Higes et al., 2007). At the physiological level, this parasite disrupts the carbohydrate (Aliferis et al., 2012; Mayack and Naug, 2009) and lipid (Li et al., 2018) metabolism of the host, which ensures the availability of nutrients for its own replication. This causes an energetic stress on infected honey bees that consume more sucrose solution than non-infected conspecifics (Alaux et al., 2010; Martín-Hernández et al., 2011; but see Aufauvre et al., 2012). *N. ceranae* also alters gene expression in the brain (Holt et al., 2013), inhibits the apoptosis of epithelial cells (Martín-Hernández et al., 2017) and deregulates immune responses (Martín-Hernandez et al., 2018). At the behavioural level, infected honey bees start foraging earlier in life (Goblirsch et al., 2013; Higes et al., 2009; Perry et al., 2015), exhibit more frequent but shorter foraging flights (Dosselli et al., 2016; Dussaubat et al., 2013; Kralj and Fuchs, 2010), show reduced homing abilities (Wolf et al., 2014) and lower olfactory learning performances (Piiroinen and Goulson, 2016; Gage et al., 2018; but see Charbonneau et al., 2016).

Over the past ten years, *N. ceranae* has been identified in an increasing number of wild bee species (bumblebees: (Graystock et al., 2013; Li et al., 2012; Plischuk et al., 2009), stingless bees: (Porrini et al. 2017)) as well as in some wasps (Porrini et al., 2017) and butterflies (Malysh et al., 2018). In the field, horizontal transmission to wild pollinators may occur through contamination of flower pollen by infected honey bees. This possibility is particularly concerning since many of these wild pollinators are solitary or present simple social organisations (Michener, 2000), and therefore lack complex social immunity behaviours that allow honey bees to collectively limit infection risks and combat parasites (Cremer et al., 2007).

The effects of *N. ceranae* on wild pollinators have been best investigated in bumblebees. Recent studies suggest that *N. ceranae* impairs the cognitive abilities of bumblebees, potentially reducing the foraging performances of entire colonies (Piiroinen et al., 2016; Piiroinen and Goulson, 2016). However, the diversity of experimental infections protocols used in these studies, and their relatively low success rates (i.e. from 0% in (Piiroinen et al., 2016) to 62% in (Graystock et al., 2013)), make it difficult to draw definitive conclusions and call for more robust standardised approaches. By contrast, 100% of infections are routinely reached in honey bee studies (Doublet et al., 2015; Roberts and Hughes, 2015).

The age of bumblebees, their level of starvation prior to exposure, and the amount of parasite spores to which they are exposed to, are all potentially important parameters that vary across current infection protocols. Highest infections levels were obtained by starving bumblebees of uncontrolled age for 8h before feeding them with 6500 spores (an empirically measured infection intensity in wild colonies) in 40% sucrose solution individually administered with a pipette (Graystock et al., 2013). Bumblebees were then maintained in smalls groups of 10 individuals with *ad libitum* access to 40% sucrose solution. Under these conditions, 62% of the bumblebees showed spores in their gut, which considerably reduced their lifespan. Exposure to higher spore dosages caused different results. Fürst et al., (2014) starved 2-days old bumblebees for 1h before feeding them with 100K spores administered in a 10 μL drop of sucrose and maintained them individually with 50% sucrose and artificial pollen *ad libitum*. Here, 34% of the bumblebees developed an infection but no effect was observed on survival (Fürst et al., 2014). Piiroinen and Goulson (2016) and Piiroinen et al., (2016) administered bumblebees of uncontrolled age 180K and 130K *N. ceranae* spores respectively with a pipette and maintained them in small groups of 10 with *ad libitum* access to 50% sucrose and pollen. In these experiments, where the bumblebees were provided high numbers of spores but starved for only 2h, 3% (Piiroinen and Goulson, 2016) or none (Piiroinen et al., 2016) of the individuals developed an infection.

Another potentially important parameter, so far unexplored, is the nutritional composition of diets provided to bumblebees after parasite exposure. Nutrition is as a key mediator of host-parasite interactions (Ponton et al., 2011) and should therefore be carefully controlled when developing standard infection procedures. Many insects increase their consumption of dietary protein in order to develop stronger immunological responses and combat parasites (Cotter et al., 2011; Lee et al., 2006; Povey et al., 2014). Increasing evidence indicate that bees given a choice of foods can carefully adjust their intake of nutrients to reach target levels maximizing fitness traits (e.g. Vaudo et al., 2016; Kraus et al., 2019; Ruedenauer et al., 2020). Although there is no direct demonstration that diet modifies the ability of bees to fight infections, bumblebees fed a low-protein diet have a reduced immune response (Brunner et al., 2014). Honey bees infected by *Nosema apis* (Rinderer and Elliott, 1977) and *N. ceranae* (Jack et al., 2016; Porrini et al., 2011; Tritschler et al., 2017) survive longer when provided protein rich pollen. Since protein consumption is needed for synthesising peptides in immune pathways (Lee et al., 2006; Mason et al., 2014; Povey et al., 2014), these results suggest that bees can adjust their nutrient intake for self-medication (Poissonnier et al., 2018).

Here we developed an experimental protocol to efficiently infect bumblebees with *N*. *ceranae* by briefly exposing individuals to controlled amounts of parasite spores and feeding them with artificial diets varying in their protein to carbohydrate ratios. We calibrated our approach by testing two spore dosages and two diets. We analysed parasite prevalence (proportion of infected bumblebees based on polymerase chain reactions, PCR), parasite intensity (spore counts in the gut) and host survival to identify best conditions for infecting bumblebees at sub-lethal doses.

## 2. Materials and Methods

### 2.1. Bees

Experiments were conducted in December 2017. Bumblebee workers (*Bombus terrestris*) of unknown age were obtained from two commercial colonies (Biobest, Belgium). Prior to the experiments, 15 workers from each colony were sampled for parasite screening (Graystock et al., 2013). The absence of *N. ceranae* and other common parasites of bumblebees (*N. bombi* and *Crithidia bombi*) in these samples was confirmed in a monoplex PCR with the primers 218MITOC (*N. ceranae:* Martín-Hernández et al., 2007), Nbombi-SSU-J (*N. bombi*: Klee et al., 2006) and CB-ITS1 (*C. bombi:* Schmid-Hempel and Tognazzo, 2010).

To validate our approach on bumblebees, Honey bees (*Apis mellifera*) were used as positive controls for parasite infection. Honey bee workers of unknown age were obtained from a *Nosema*-free (*N. ceranae* and *N. apis*) colony maintained at our experimental apiary (University Paul Sabatier, Toulouse). The absence of parasite in the colony was confirmed in a duplex PCR with the primers 218MITOC and 321APIS on a sample of 15 workers (Martín-Hernández et al., 2007).

### 2.2. Parasites

*N. ceranae* spores were obtained from a naturally infected honey bee colony. The abdomens of 15 honey bee foragers were crushed into dH_2_O, and the homogenate was examined using light microscopy (x400). Samples showing spores were checked in a duplex PCR (Martín-Hernández et al., 2007) to verify for the presence of *N. ceranae* and the absence of *N. apis* (spores of both parasites have similar morphologies; Fries et al., (2006)). Positive honey bee gut homogenates were used to prepare the spore solutions for the experimental infections, following a standard purification protocol (Fries et al., 2013). Spore solutions were prepared no more than one week before the infections. Briefly, 1 mL of the honey bee gut homogenate was centrifuged at 5000 rpm for 5 min. The supernatant containing tissue debris was discarded and the pellet containing the spores was re-suspended in 0.5 mL of dH_2_O by vortexing. The sample was centrifuged and washed into dH_2_O two times more to obtain a spore solution of 85% purity (Fries et al., 2013). *N. ceranae* spores were counted using an improved Neubauer haemocytometer (Cantwell, 1970) in a light microscope (x400). Counts were made in five squares of 0.2 × 0.2 mm^2^ area in both chambers of the haemocytometer and the total number of spores was averaged. The final inoculum concentration was adjusted to either 7500 spores/μL or 15000 spores/μL in 20% (v/v) of sucrose solution. All the inocula were prepared the day of the experimental infections.

### 2.3. Infections

Bumblebees and honey bees were exposed to solutions of *N. ceranae* spores on the same day. Based on preliminary observations showing that different starvation durations are needed to elicit feeding in bumblebees and honey bees, bumblebees and honey bees were manipulated separately. At 9.00 am (GMT+1), 300 bumblebees were isolated in empty Petri Dishes (20 mm <u>Ø</u>) and starved for 5h. At 12.00 pm, 300 honey bees were isolated in the same conditions but for 2h only. At 2.00 pm, bumblebees were individually exposed to a drop of 20 μL of sucrose solution (20% v/v) containing either 0 spore (control), 150K spores or 300K spores. Honey bees were exposed to a drop of sucrose solution containing either 0 spore (control) or 150K spores. The sucrose was delivered with a micropipette in the Petri dish of each bee. Only the bees that consumed the entire drop within 2 two hours were kept for the experiments (272 out of the 300 bumblebees, 258 out of 300 honey bees, see details in supplementary Table 1).

### 2.4. Artificial diets

Immediately after parasite exposure, bees were allocated to one of two artificial diets, which generated 10 experimental groups. The number of individuals in each group varied between 44 and 50 for bumblebees, and 47 and 77 for honey bees (supplementary Table 1). Diets were liquid solutions containing either a low protein to carbohydrate ratio (P:C 1:150, hereafter “low-protein diet”) or a high protein to carbohydrate ratio (P:C 1:5, hereafter “high-protein diet”). The two diets were prepared with a fixed total amount of nutrients (P + C content of 170 g/L). Carbohydrates were supplied as sucrose (Euromedex, France). Proteins consisted in a mixture of casein and whey (4:1) (Nutrimuscle, Belgium). Both diets contained 0.5% of vitamin mixture for insects (Sigma, Germany).

### 2.5. Survival

Bumblebees and honey bees were individually kept in a Petri dish with a hole (1 cm Ø) on the top lid, in which was placed a gravity feeder containing one of the two diets. The feeder consisted in a 1.5 mL Eppendorf tube with a hole at its basis through which the bees could insert their proboscis and ingest liquid food. All bees were kept in two identical incubators (Pol-Eko, Poland) at 26°C, with a photoperiod of 12h light: 12h dark, for 21 days. Each incubator contained the same proportion of bees from the 10 experimental groups (supplementary Table 1). Every day, diets were renewed and dead bees were stored at −20 °C for molecular analyses. Bees that survived the entire experiment were freeze-killed on day 21.

### 2.6. Prevalence and intensity of *N. ceranae*

Parasite prevalence (presence in the gut of bees) was assessed in bumblebees and honey using PCR (see example Figure 1A). The gut of each bee was extracted and homogenised in dH_2_O and vortexed with 2 mm glass beads (Labbox Labware, Spain). Genomic DNA was extracted using Proteinase K (20 mg/mL; Euromedex, France) and 1 mM of Tris-EDTA Buffer (pH = 8). PCR conditions were adapted from Martín-Hernández et al., (2007) for a monoplex PCR with the 218MITOC primers specific for *N. ceranae*. PCR reactions were carried out in 48-well microtitre plates in a S1000™ Thermal Cycler (Biorad, CA). Each reaction contained 1.5 U of Taq Polymerase (5 U/μL; MP Biomedicals, CA), 1x PCR Direct Loading Buffer (MP Biomedicals, CA), 0.4 μM of each pair of primers (Martín-Hernández et al., 2007), 200 μM of dNTPs (Jena Biosciences, Germany), 0.48 μg/μL of BSA (Sigma, Germany) and 2.5 μL of DNA sample in a final volume of 25 μL. Thermal conditions consisted in an initial denaturing step of 94 °C for 2 min, followed by 35 cycles of 94 °C for 30 s, 61.8°C for 45 s and 72 °C for 2 min, with a final elongation step of 72 °C for 7 min. The length of PCR products (i.e. 218 pb) was checked in a 1.2% agarose gel electrophoresis stained with SYBR Safe DNA Stain (Edvotek, Washington DC).

**Figure 1.**
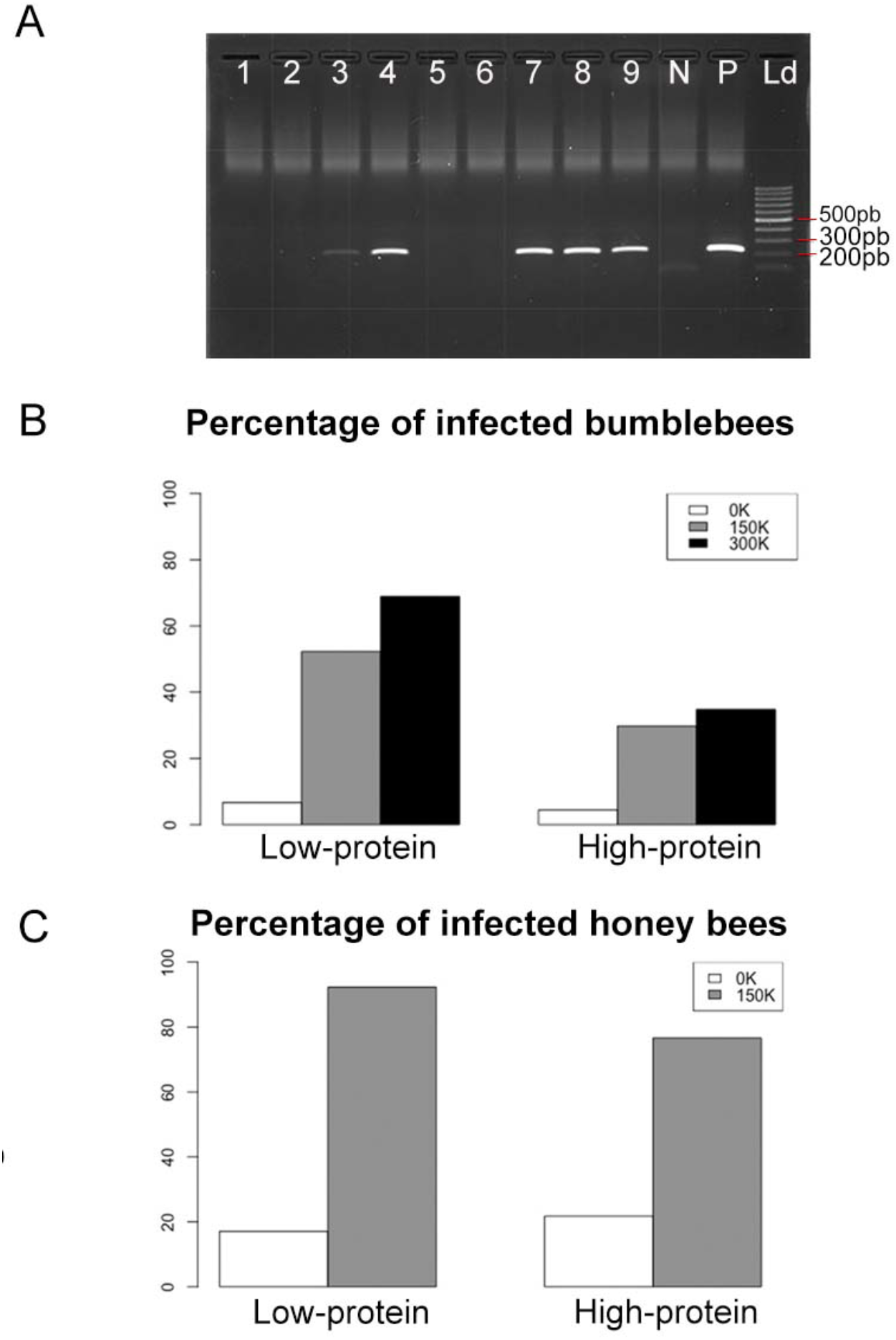
Parasite prevalence. (A) Agarose electrophoresis gel with negative (lanes 1, 2, 5 and 6) and positive (lanes 3, 4, 7-9) bumblebees to *N. ceranae* determined by PCR. N: negative control of PCR; P: positive control of PCR; Ld: molecular size marker (100 pb). Percentage of (B) bumblebees and (C) honey bees infected by *N. ceranae* (PCR positive) for all diets and spore dosages. Bumblebee groups contained between 44 and 50 individuals. Honey bee groups contained between 47 and 77 individuals (see details in supplementary Table 1).

Parasite intensity (spore load in the gut) was estimated in positive PCR bumblebee samples under a light microscope (x400). The number of spores was counted as described above (see Section 1.2 and example Figure 2A). Random samples of *Nosema*-negative PCR bumblebees were also screened to confirm the absence of parasite spores.

**Figure 2.**
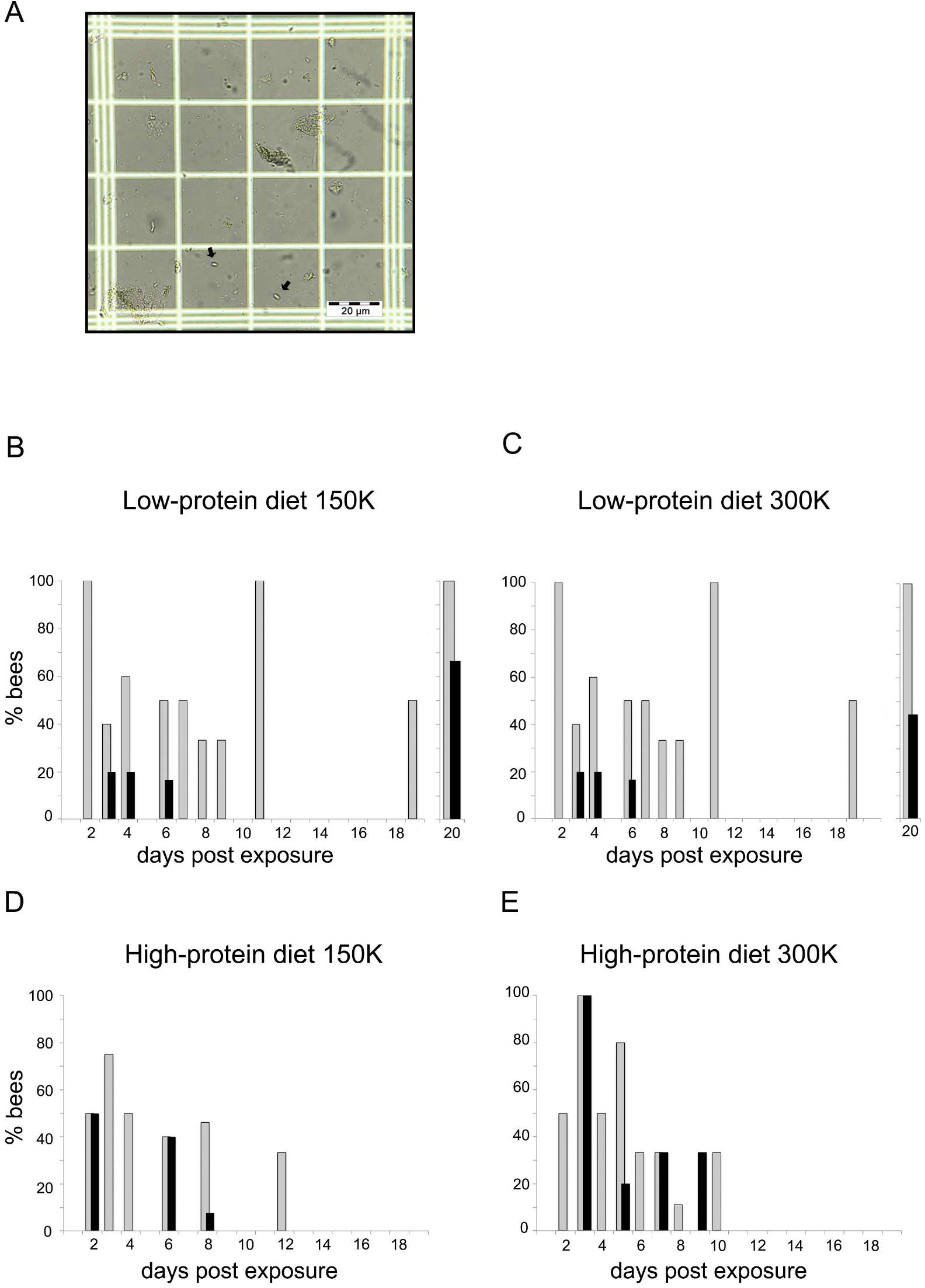
Parasite loads. (A) *N. ceranae* spores observed in an infected bumblebee (optical microscope, x 400). (B-E) Proportion of dead bumblebees at different days post exposure that were positive to *N. ceranae* in a PCR (grey bars) and showed spores in their gut (black bars), for both diets (high- and low-protein diets) and spore dosages (150K and 300K). Each group contained between 44 and 50 individuals (see details in supplementary Table 1).

### 2.7. Statistical analyses

Bumblebees were analysed for parasite prevalence, parasite intensity and survival. Honey bees were used as positive control for parasite infection, and were therefore only analysed for parasite prevalence. All analyses were performed in R v. 1.0.143 (R Development Core Team). All means are shown with standard errors (mean ± S.E).

The effects of diet, spore dosage and their interaction on parasite prevalence (proportion of individuals infected after parasite exposure) were tested using Generalized Linear Models with a binomial error structure (binomial GLMs). The effect of diet, spore dosage and their interaction on the proportion of bees showing *N. ceranae* spores in their gut were tested using a binomial GLM. The effect of diet, spore dosage and their interaction on parasite intensity (number of spores per bee) were tested using a negative binomial GLM (because of the large over-dispersion of the data). The parasite loads of individuals bees that survived until the end of the experiment (day 21) and those that died before were compared using a negative binomial GLM with a binary predictor (e.g. dead or alive at day 21) as fixed effect. GLMs with binomial distribution errors were fitted using the *glm* function in the R package “stats”. GLMs with a negative binomial error distribution were fitted using the *glm.nb* function in the R package “MASS” (Venables and Ripley, 2002). All the models were tested for interactions among predictors (e.g. diet, dosage and PCR results). Interactions that did not improve the fitting were removed from the models using the Akaike information criterion (AIC) for model comparison (Akaike, 1985).

The survival of bumblebees was analysed using a Kaplan-Meier test curve with the function *survfit* in the R package “Survival” (Therneau and Grambsch, 2000). The effects of spore dosage, diet, infectious status and their interactions were analysed using Cox proportional-hazards regression models (function *coxph* in the R package “Survival”), followed by a Tukey post-hoc test to account for pairwise comparisons. In all models, colony origin and incubator identity were included as random factors.

## 3. Results

### 3.1. Parasite prevalence

To assess infection rates, bumblebees and honey bees were analysed with PCR for the presence of parasites after the 21 days of the experiment. In total, 46.15% of the bumblebees (84 out of 182, Figure 1B and Table 1) and 83.8% of the honey bees (119 out of 142, Figure 1C) exposed to *N. ceranae* were PCR positive, which validates our protocol. Note however that a proportion of control (non-exposed) individuals (5.5% of bumblebees, 19.8% honey bees) were found PCR positive (Figures 1B and 1C). This suggests that our colonies were not entirely free of parasites prior to the experiments or that horizontal cross-contamination occurred during the experiments. All these infected control bees were excluded from further analyses.

**Table 1.** Number of bumblebees exposed to *N. ceranae*, infected with parasite (PCR positive), and showing spores in their gut. The percentage of bumblebees infected relative to those exposed, and the percentage of bumblebees showing spores in their gut relative to the number of PCR positive individuals are given into brackets.

| Diet | Spore dosage | Exposed | Infected | Showing spores |
| --- | --- | --- | --- | --- |
| Low-protein | 150K | 44 | 23 (52.27%) | 7 (30.4%) |
|  | 300K | 45 | 31 (68.88%) | 9 (29.03%) |
| High-protein | 150K | 47 | 14 (29.78%) | 4 (28.5%) |
|  | 300K | 46 | 16 (34.78%) | 6 (37.5%) |
| Total |  | 182 | 84 (46.15%) | 26 (30.9%) |

Diets significantly influenced the proportion of bees that became infected after exposure to the parasite (Figures 1B and 1C; Table 1). A significantly larger number of bumblebees became infected when fed the low-protein diet than when fed the high-protein diet (GLM_binomial_: Estimate = 1.186 (± 0.313), z = 3.79, *p* < 0.001), irrespective of spore dosage (GLM_binomial_: Estimate = 0.4870 (± 0.313), z = 1.503, p = 0.133). Likewise, a larger number of honey bees were infected when fed the low-protein than when fed the high protein-diet (GLM_binomial_: Estimate = 1.298 (± 0.538), z = 2.41, *p* = 0.016). Therefore, feeding a low-protein diet induced highest infection rates in both species. Interestingly all the bumblebees that were fed the low-protein diet and that survived until day 21 were PCR positive to *N. ceranae* (6 bumblebees for 150K, 9 bumblebees for 300K). By contrast, the only bumblebee that was fed the high-protein diet and that survived until day 21 (300K) was not infected.

### 3.2. Parasite loads

To assess the intensity of the infection, the gut homogenates of dead PCR positive bumblebees were screened under microscope for the presence of *N. ceranae* spores.

Spores were observed in bumblebee guts for both diets (Figures 2B-E; see supplementary Table 1 for details). Comparing the presence of spores in bumblebees that did not die on the same day yield some information about the dynamics of spore production. For the low-protein diet, spores were seen from day 3 to day 20 post exposure with 150K spores (Figure 2B) and from day 4 to day 20 post exposure with 300K spores (Figure 2C). Spores were never observed between day 8 and day 15. This may be due to the low mortality rate of bumblebees on these days. For the high-protein diet, spores were seen from day 2 to 8 post exposure with 150K spores (Figure 2D), and from day 3 to day 9 post exposure with 300K spores (Figure 2E).

Overall 30.9% (26 out of 84) of the infected bumblebees showed *N. ceranae* spores in their gut (Table 1). This estimation of prevalence based on spore loads is lower than that based on PCR screening (Figure 1B and Table 1) because, here, the intracellular stages of the parasite (e.g. meronts, sporonts) cannot be detected. The proportion of bumblebees with spores in their gut was similar irrespective of the diet (GLM_binomial___diet_: Estimate = 0.176 (± 0.489), z = 0.36, *p* = 0.718) and the spore dosage (GLM_binomial___spore dosage_: Estimate = 0.109 (± 0.477), z = 0.23, *p* = 0.819). Spore loads were highly variable across individuals, with an average of 367.31 (± 72.03) spores per bumblebee. The spore dosage the bees were exposed to had no effect on the spore loads found in their gut (GLM_negative-binomial_: Estimate = −0.002 (± 0.002), z = −0.73, *p* = 0.464). On the contrary, the diet had a significant effect on spore load (GLM_negative-binomial_: Estimate = 1.294 (± 0.347), z = 3.73, *p* < 0.001), leading to higher amounts of spores in bumblebees fed the high-protein diet (662.50 ± 222.71 spores per bee) than in bumblebees fed the low-protein diet (182.81 ± 40.99 spores per bee).

Considering bumblebees that were fed the low-protein diet and that survived until day 20 post exposure, only 4 out of 6 individuals exposed to 150K spores (175 ± 87.5 spores per bee; Figure 2B) and 4 out 9 individuals exposed to 300K spores (125 ± 62.5 spores per bee; Figure 2C) showed spores in their gut. These bumblebees showed similar spore loads than those fed the same diet but that died before day 20 post exposure (GLM_negative-binomial:_ Estimate = −0.363 (± 0.389), z = −0.93, *p* = 0.350).

### 3.3. Survival

To test the effect of spore dosage and diet on the physiology of bumblebees, survival was analysed during the 20 days after parasite exposure. Here the infection status of each bumblebee was based on the PCR results.

Overall, bumblebees fed the high-protein diet had lower survival than bumblebees fed the low-protein diet (Figure 3; Cox: estimate = − 0.75 (± 0.14), z = −5.17, p < 0.001). Bumblebees fed the high-protein diet (Figure 3A) showed similar mortality rate irrespective of spore dosage (Table 2) and infectious status (supplementary Table 2). By contrast, bumblebees fed the low-protein diet (Figure 3B) had a significantly reduced survival when exposed to spores than when non-exposed (Table 2). For this diet, 50% of exposed bumblebees died between days 6 and 7 post exposure. This lethal time 50 (LT_50_) was not reached for the non-exposed (control) bumblebees. Interestingly, in the low-protein diet, infected bumblebees tended to die faster than control bumblebees, but survived significantly longer than bumblebees exposed and non-infected (Figure 3B; supplementary Table 2). Therefore, both parasite exposure and infection had negative effects on the survival of bumblebees maintained on the low-protein diet.

**Table 2.**
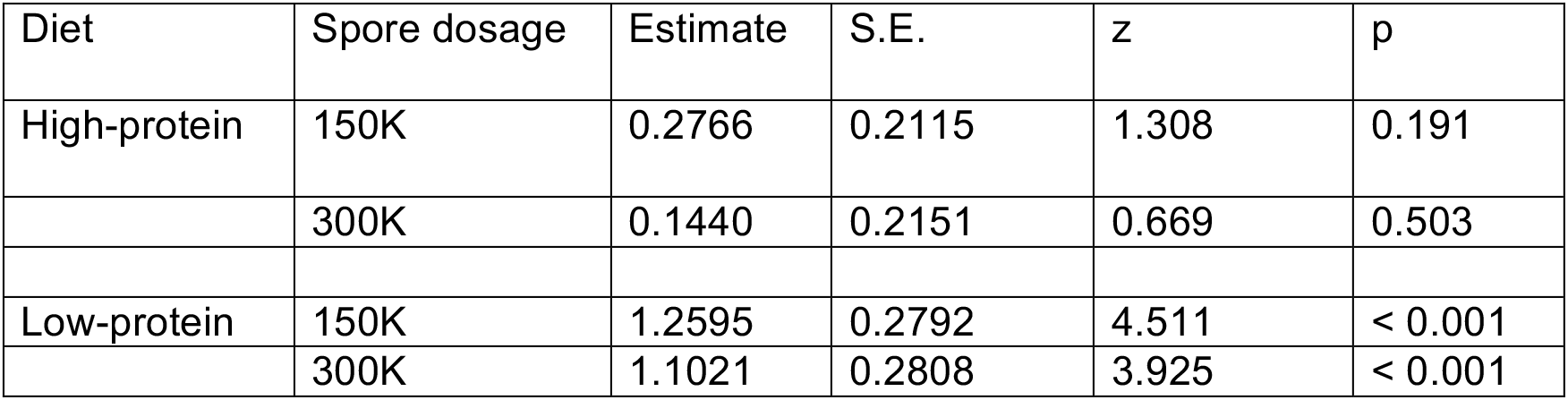
Cox Regression Model testing the effect of spore dosage on the survival of bumblebees for each diet.

| Diet | Spore dosage | Estimate | S.E. | z | p |
| --- | --- | --- | --- | --- | --- |
| High-protein | 150K | 0.2766 | 0.2115 | 1.308 | 0.191 |
|  | 300K | 0.1440 | 0.2151 | 0.669 | 0.503 |
| Low-protein | 150K | 1.2595 | 0.2792 | 4.511 | < 0.001 |
|  | 300K | 1.1021 | 0.2808 | 3.925 | < 0.001 |

**Figure 3.**
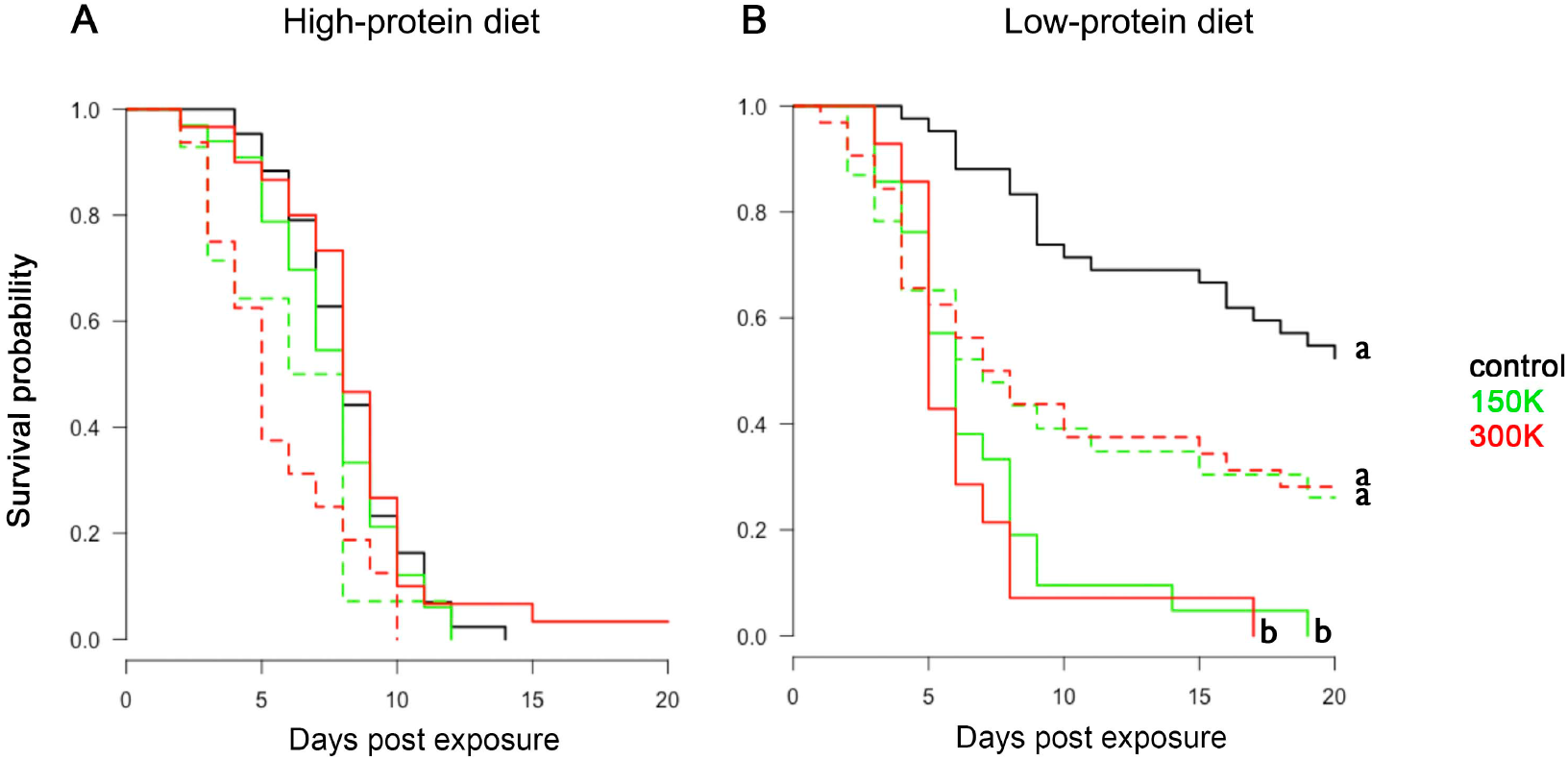
Survival analyses. Survival probability of bumblebees fed the high-protein diet (A) and the low-protein diet (B), for the different spore dosages (control, 150K and 300K) across time. Infected (dashed lines) and non-infected (solid lines) bumblebees are differentiated. Different letters associated to curves indicate statistical differences in survival (p<0.05; post-hoc Tukey test after Cox-regression model in a Kaplan Meier test curve, see supplementary Table 3). Each group contained between 44 and 50 individuals (see details in supplementary Table 1).

## 4. Discussion

Several recent studies suggest that *N. ceranae* negatively affect bee behaviour and cognition, which may have dramatic consequences for colony growth and survival (Gage et al., 2018; Piiroinen and Goulson, 2016). In bumblebees, these results remain difficult to interpret due to the low efficiency of experimental infection protocols. Here we exploited recent insight into the nutritional ecology of bees (e.g. Vaudo et al., 2016; Kraus et al., 2019; Ruedenauer et al., 2020) to develop a reliable method to infect bumblebees with *N. ceranae*. We identified key effects of diet composition on bumblebee infection rates and survival, indicating a complex interaction between diet, parasite development and host health.

Overall 46% of the bumblebees exposed to *N. ceranae* spores were infected, a percentage that increased up to 70% when only considering bumblebees that were exposed to 300K spores and maintained on a low-protein high-carbohydrate diet. These levels of prevalence are similar to those obtained by Graystock et al., (2013) (62% infection rate) who used a 20 to 45 times lower infection dose (measured in naturally infected colonies: 6500 spores/bee) administered with a micropipette directly into the bee mouth parts. These rates of successful infections are higher than all previous studies where bumblebees were exposed to an order of 10^5^ spores/bee either delivered as a drop of solution in a Petri Dish (39%; Fürst et al., 2014) or with a micropipette directly into the bee mouth (0-3%; Piiroinen and Goulson, 2016; Piiroinen et al., 2016).

We did not find any clear effect of parasite dosage (150K or 300K spores per bee) on prevalence and survival. Comparing our results to that of previous studies does not indicate such effect neither, which suggests that above a certain concentration of parasite spores (e.g. 6500 spores/bee in Graystock et al. 2013), parasite dosage is not an important factor determining the success of an infection. By contrast, we found a strong effect of diet on parasite prevalence and spore loads. Previous studies indicate that pollen intake affects honey bee physiology and tolerance to *N. ceranae* (Basualdo et al., 2014; Di Pasquale et al., 2013). Using artificial diets with controlled amounts of protein and carbohydrates, we demonstrate that food macronutrient balance substantially influences both the infection rate and the survival of bumblebees. A higher number of bumblebees were infected when fed the low-protein high-carbohydrate diet, suggesting that different blends of nutrients differently affect the germination of the parasite (but see Jack et al., 2016). The digestion of the different macronutrients in the bumblebee gut, whose microbiota is mostly composed by sugar-fermentative bacteria (Mohr and Tebbe, 2006), may lead to changes in the gut environment that could favour the parasite germination, such as shifts in pH (Wittner and Weiss, 1999). Furthermore, because *N. ceranae* exploits the metabolism of carbohydrates to obtain energy from its host (Mayack and Naug, 2009) a higher carbohydrate consumption by infected bumblebees may benefit the establishment of *N. ceranae*. Alike previous studies on honey bees that showed that the supplement of pollen in diet increase the parasite intensity (Basualdo et al., 2014; Fleming et al., 2015; Jack et al., 2016), we found that bumblebees with access to higher amounts of protein show higher spore loads in their gut, suggesting that proteins benefit the replication of the parasite.

The effect of diet was also evident on bumblebee survival. *N. ceranae* reduced the survival of bumblebees, specifically in the low-protein high-carbohydrate diet, where mortality rate increased by day 3 post exposure. This result is consistent with the time described for *N. ceranae* to produce new spores in the honey bee gut (Higes et al., 2007). Similarly to Graystock et al., (2013), we found that most of the infected bumblebees (67.5%) died without showing spores in their gut, suggesting that the parasite was not able to complete its cycle at a detectable level (i.e. produce new spores). The lack of spores in the bumblebee guts could be due to a migration of the parasite to the fat body of their host where it cannot reproduce (Graystock et al., 2013). Alternately, this could be a consequence of a high virulence of the parasite. Emergent parasites can be very virulent for a novel host causing its death before reproducing and releasing their offspring (Schmid-Hempel, 1998). Although it is not clear when *N. ceranae* started to colonise bumblebee hosts (including *B. terrestris*), the parasite was first identified in bumblebees in 2005 (Plischuk et al., 2009), implying a short coevolution time in which the parasite may not have had time to fully adapt its virulence to its diverse host species. Interestingly, the absence of parasite spores in the gut of bumblebees that died at different times could reflect the ability of bumblebees to clear the parasite. The fact that infected bumblebees fed the low-protein high-carbohydrate diet survived longer than exposed non-infected bumblebees fed the same diet suggests that the mechanisms allowing parasite clearance incur a higher metabolic cost, here manifested as a reduction of bumblebee lifespan, than parasite tolerance. Nonetheless, given that only dead bumblebees were checked for spores and that spore loads are generally higher in live bees than in dead ones (Zheng et al., 2014), studying the dynamics of *N. ceranae* spore production in live bumblebees (for instance by analysing faeces) will be necessary to determine whether these insects are able to get rid of the parasite.

The fact that bumblebees showed low parasite prevalence but died faster when fed a high-protein low-carbohydrate diet suggests that immunity and lifespan are maximized at different nutritional balances. Several studies show how an animals’ diet can differently influence the expression of key life-history traits, forcing animals to trade-off between optimizing multiple traits simultaneously (e.g. Bunning et al., 2016, 2015; Rapkin et al., 2018). In insects, lifespan is typically enhanced on high-carbohydrate diets whereas reproduction is maximized on high-protein diets (fruit flies: Lee et al., 2008; Fanson et al., 2009; Reddiex et al., 2013; Jensen et al., 2015; Semaniuk et al., 2018, crickets: Maklakov et al., 2008). Immunity and reproduction also display differences in nutritional requirements (fruit flies: Ponton et al., 2011, 2015; decorated crickets: Rapkin et al., 2018; leafworm: Cotter et al., 2011). Honey bees survive longer on high-carbohydrate diets (Altaye et al., 2010) and pollen (the main source of protein) favours the survival of individuals infected with *N. ceranae* (Basualdo et al., 2014; Fleming et al., 2015; Jack et al., 2016; Zheng et al., 2014). Whether sterile bumblebee workers trade-off between over-ingesting protein and under-ingestion of carbohydrates in order to reduce parasite establishment at the expense of a shorter lifespan is an open question. In these social insects, such individual strategy may considerably reduce the risks of contamination of other workers within the colony (Poissonnier et al., 2018).

Developing standard protocols to characterize the sublethal effects of parasites and pathogens on bees has become a major challenge for understanding bee population declines (Gómez-Moracho et al., 2017). Here we demonstrated that diet is key in determining infection rates of bumblebees exposed to *N. ceranae*. Highest infection rates and longest survivals were obtained with a high-carbohydrate low-protein diet, thereby providing ideal conditions for investigating potential effects of *N. ceranae* on bee behaviour and cognition. Other parameters, not tested here, may also be of importance and investigated in future studies. For instance, both our study and that of Graystock et al. (2013) starved bumblebees for at least 5 h prior to parasite exposure, which may have increased the probability of parasite establishment. Variations in the age of the bumblebee tested may also explain some of these differences. In the only study that controlled for age, Fürst et al., (2014) infected 2-days old bumblebees (post eclosion from the pupa) and obtained a lower infection rate than that presented here and no effect on bumblebee mortality, suggesting that young bumblebees are less susceptible to the parasite. Finally, we cannot exclude differences in the virulence of the *N. ceranae* strains used for experimental infection across studies (but see Van der Zee et al., 2014). We hope that our infection protocol will facilitate further studies on the interactions between *N. ceranae* and bumblebees in order to better assess the risks this emergent parasite represents for wild pollinators.

## Acknowledgements

We thank Pascale Belenguer, Lucie Hotier, Audrey Dussutour and Enikö Csata for their help and advice.

## Funding

TGM was funded by a postdoctoral fellowship from the Fyssen Foundation. PH was funded by the ‘Laboratoire d’Excellence (LABEX)’ TULIP (ANR-10-LABX-41). TD, CP and ML were funded by a grant from the Agence Nationale de la Recherche to ML (ANR-16-CE02-0002-01).

## Notes

#### Summary of Updates

Parasites alter the physiology and behaviour of their hosts. In domestic honey bees, the microsporidia Nosema ceranae induces an energetic stress and impairs the behaviour of foragers, potentially leading to colony collapse. Whether this emerging parasite similarly affects wild pollinators is little understood because of the low success rates of experimental infection protocols. Here we established a new apporach for infecting bumblebees (Bombus terrestris) with controlled amounts of N. ceranae, by briefly exposing individual bumblebees to a sucrose solution containing parasite spores, before feeding them with artificial diets. We validated our protocol by testing the effect of two spore dosages and two diets varying in their protein to carbohydrate ratio, on the prevalence of the parasite (proportion of infected bees), the intensity of infection (spore count in the gut), and the survival of bumblebees. Insects fed a low-protein high-carbohydrate diet showed highest parasite prevalence (up to 70%) but lived longest, suggesting that immunity and survival of bumblebees are maximised at different protein to carbohydrate ratios. Spore dosage had no effect on parasite infection rate and host survival. The identification of experimental conditions for successfully infecting bumblebees with N. ceranae in the lab will facilitate investigations of the sub-lethal effects of this parasite on the behaviour and cognition of wild pollinators.

## References

Akaike, H., 1985. Prediction and Entropy, in: A celebration of statistics. Springer New York, New York, NY, pp. 1–24. doi:10.1007/978-1-4613-8560-8_1

Alaux, C., Brunet, J.-L., Dussaubat, C., Mondet, F., Tchamitchan, S., Cousin, M., Brillard, J., Baldy, A., Belzunces, L.P., Le Conte, Y., 2010. Interactions between *Nosema* microspores and a neonicotinoid weaken honeybees (*Apis mellifera*). Environ. Microbiol. 12, 774–82. doi:10.1111/j.1462-2920.2009.02123.x

Aliferis, K., Copley, T., Jabaji, S., 2012. Gas chromatography-mass spectrometry metabolite profiling of worker honey bee (*Apis mellifera L*.) hemolymph for the study of *Nosema ceranae* infection. J. Insect Physiol. 58, 1349–59. doi:10.1016/j.jinsphys.2012.07.010

Altaye, S.Z., Pirk, C.W.W., Crewe, R.M., Nicolson, S.W., 2010. Convergence of carbohydrate-biased intake targets in caged worker honeybees fed different protein sources. J. Exp. Biol. 213, 3311–3318. doi:10.1242/jeb.046953

Antúnez, K., Martín-Hernández, R., Prieto, L., Meana, A., Zunino, P., Higes, M., 2009. Immune suppression in the honey bee (*Apis mellifera*) following infection by *Nosema ceranae* (Microsporidia). Environ. Microbiol. 11, 2284–2290. doi:10.1111/j.1462-2920.2009.01953.x

Aufauvre, J., Biron, D.G., Vidau, C., Fontbonne, R., Roudel, M., Diogon, M., Viguès, B., Belzunces, L.P., Delbac, F., Blot, N., 2012. Parasite-insecticide interactions: a case study of *Nosema ceranae* and fipronil synergy on honeybee. Sci. Rep. 2, 326. doi:10.1038/srep00326

Basualdo, M., Barragán, S., Antúnez, K., 2014. Bee bread increases honeybee haemolymph protein and promote better survival despite of causing higher *Nosema ceranae* abundance in honeybees. Environ. Microbiol. Rep. 6, 396–400. doi:10.1111/1758-2229.12169

Brunner, F.S., Schmid-Hempel, P., Barribeau, S.M., 2014. Protein-poor diet reduces host-specific immune gene expression in *Bombus terrestris*. Proc. R. Soc. B Biol. Sci. 281, 20140128–20140128. doi:10.1098/rspb.2014.0128

Bunning, H., Bassett, L., Clowser, C., Rapkin, J., Jensen, K., House, C.M., Archer, C.R., Hunt, J., 2016. Dietary choice for a balanced nutrient intake increases the mean and reduces the variance in the reproductive performance of male and female cockroaches. Ecol. Evol. 6, 4711–4730. doi:10.1002/ece3.2243

Bunning, H., Rapkin, J., Belcher, L., Archer, C.R., Jensen, K., Hunt, J., 2015. Protein and carbohydrate intake influence sperm number and fertility in male cockroaches, but not sperm viability. Proc. R. Soc. B Biol. Sci. 282. doi:10.1098/rspb.2014.2144

Cantwell G.E. 1970. Standard methods for counting *Nosema* spores, Am. Bee J. 110, 222–223.

Cotter, S.C., Simpson, S.J., Raubenheimer, D., Wilson, K., 2011. Macronutrient balance mediates trade-offs between immune function and life history traits. Funct. Ecol. 25, 186–198. doi:10.1111/j.1365-2435.2010.01766.x

Cremer, S., Armitage, S.A.O., Schmid-Hempel, P., 2007. Social Immunity. Curr. Biol. 17, R693–R702. doi:10.1016/j.cub.2007.06.008

Di Pasquale, G., Salignon, M., Le Conte, Y., Belzunces, L.P., Decourtye, A., Kretzschmar, A., Suchail, S., Brunet, J.-L., Alaux, C., 2013. Influence of pollen nutrition on honey bee health: do pollen quality and diversity matter? PLoS One 8, e72016. doi:10.1371/journal.pone.0072016

Dosselli, R., Grassl, J., Carson, A., Simmons, L.W., Baer, B., 2016. Flight behaviour of honey bee (*Apis mellifera*) workers is altered by initial infections of the fungal parasite *Nosema apis.* Sci. Rep. 6, 36649. doi:10.1038/srep36649

Doublet, V., Natsopoulou, M.E., Zschiesche, L., Paxton, R.J., 2015. Within-host competition among the honey bees pathogens *Nosema ceranae* and Deformed wing virus is asymmetric and to the disadvantage of the virus. J. Invertebr. Pathol. 124, 31–34. doi:10.1016/j.jip.2014.10.007

Dussaubat, C., Maisonnasse, A., Crauser, D., Beslay, D., Costagliola, G., Soubeyrand, S., Kretzchmar, A., Le Conte, Y., 2013. Flight behavior and pheromone changes associated to *Nosema ceranae* infection of honey bee workers (*Apis mellifera*) in field conditions. J. Invertebr. Pathol. 113, 42–51. doi:10.1016/j.jip.2013.01.002

Fanson, B.G., Weldon, C.W., Pérez-Staples, D., Simpson, S.J., Taylor, P.W., 2009. Nutrients, not caloric restriction, extend lifespan in Queensland fruit flies (*Bactrocera tryoni*). Aging Cell 8, 514–523. doi:10.1111/j.1474-9726.2009.00497.x

Fleming, J.C., Schmehl, D.R., Ellis, J.D., 2015. Characterizing the impact of commercial pollen substitute diets on the level of *Nosema* spp. in honey bees (*Apis mellifera L*.). PLoS One 10. doi:10.1371/journal.pone.0132014

Fries, I., Chauzat, M.-P., Chen, Y.-P., Doublet, V., Genersch, E., Gisder, S., Higes, M., McMahon, D.P., Martín-Hernández, R., Natsopoulou, M., Paxton, R.J., Tanner, G., Webster, T.C., Williams, G.R., 2013. Standard methods for *Nosema* research. J. Apic. Res. 52, 1–28. doi:10.3896/IBRA.1.52.1.14

Fries, I., Martín, R., Meana, A., García-Palencia, P., Higes, M., 2006. Natural infections of *Nosema ceranae* in European honey bees. J. Apic. Res. 47, 230–233. doi:10.3896/IBRA.1.45.4.13

Fürst, M.A., McMahon, D.P., Osborne, J.L., Paxton, R.J., Brown, M.J.F., 2014. Disease associations between honeybees and bumblebees as a threat to wild pollinators. Nature 506, 364–366. doi:10.1038/nature12977

Gage, S.L., Kramer, C., Calle, S., Carroll, M., Heien, M., DeGrandi-Hoffman, G., 2018. *Nosema ceranae* parasitism impacts olfactory learning and memory and neurochemistry in honey bees (*Apis mellifera*). J. Exp. Biol. 221, jeb161489. doi:10.1242/jeb.161489

García-Palencia, P., Martín-Hernández, R., González-Porto, A.-V., Marin, P., Meana, A., Higes, M., 2010. Natural infection by *Nosema ceranae* causes similar lesions as in experimentally infected caged-workers honey bees (*Apis mellifera*). J. Apic. Res. 49, 278–283. doi:10.3896/IBRA.1.49.3.08

Goblirsch, M., Huang, Z.Y., Spivak, M., 2013. Physiological and behavioral changes in honey bees (*Apis mellifera*) induced by *Nosema ceranae* infection. PLoS One 8, e58165. doi:10.1371/journal.pone.0058165

Gómez-Moracho, T., Heeb, P., Lihoreau, M., 2017. Effects of parasites and pathogens on bee cognition. Ecol. Entomol. 42, 51–64. doi:10.1111/een.12434

Graystock, P., Yates, K., Darvill, B., Goulson, D., Hughes, W.O.H., 2013. Emerging dangers: deadly effects of an emergent parasite in a new pollinator host. J. Invertebr. Pathol. 114, 114–119. doi:10.1016/j.jip.2013.06.005

Higes, M., García-Palencia, P., Martín-Hernández, R., Meana, A., 2007. Experimental infection of *Apis mellifera* honeybees with *Nosema ceranae* (Microsporidia). J. Invertebr. Pathol. 94, 211–7. doi:10.1016/j.jip.2006.11.001

Higes, M., Martín-Hernández, R., Garrido-Bailón, E., González-Porto, A. V, García-Palencia, P., Meana, A., Del Nozal, M.J., Mayo, R., Bernal, J.L., 2009. Honeybee colony collapse due to *Nosema ceranae* in professional apiaries. Environ. Microbiol. Rep. 1, 110–113. doi:10.1111/j.1758-2229.2009.00014.x

Holt, H.L., Aronstein, K. a, Grozinger, C.M., 2013. Chronic parasitization by *Nosema* microsporidia causes global expression changes in core nutritional, metabolic and behavioral pathways in honey bee workers (*Apis mellifera*). BMC Genomics 14, 799. doi:10.1186/1471-2164-14-799

Jack, C.J., Uppala, S.S., Lucas, H.M., Sagili, R.R., 2016. Effects of pollen dilution on infection of *Nosema ceranae* in honey bees. J. Insect Physiol. 87, 12–19. doi:10.1016/j.jinsphys.2016.01.004

Jensen, K., McClure, C., Priest, N.K., Hunt, J., 2015. Sex-specific effects of protein and carbohydrate intake on reproduction but not lifespan in *Drosophila melanogaster*. Aging Cell 14, 605–615. doi:10.1111/acel.12333

Klee, J., Tek Tay, W., Paxton, R.J., 2006. Specific and sensitive detection of *Nosema bombi* (Microsporidia: Nosematidae) in bumble bees (*Bombus* spp.; Hymenoptera: Apidae) by PCR of partial rRNA gene sequences. J. Invertebr. Pathol. 91, 98–104. doi:10.1016/j.jip.2005.10.012

Klein, S., Cabirol, A., Devaud, J.-M., Barron, A.B., Lihoreau, M., 2017. Why bees are so vulnerable to environmental stressors. Trends Ecol. Evol. doi:10.1016/j.tree.2016.12.009

Kralj, J., Fuchs, S., 2010. *Nosema* sp. influences flight behavior of infected honey bee (*Apis mellifera*) foragers. Apidologie 41, 21–28. doi:10.1051/apido/2009046

Kraus, S., Gomez-Moracho, T., Pasquaretta, C., Latil, G., Dussutour, A., Lihoreau, M., 2019. Bumblebees adjust protein and lipid collection rules to the presence of brood. Curr Zool. 65, 437–446. doi:10.1093/cz/zoz026

Lee, K.P., Cory, J.S., Wilson, K., Raubenheimer, D., Simpson, S.J., 2006. Flexible diet choice offsets protein costs of pathogen resistance in a caterpillar. Proc. R. Soc. B. Biol. Sci. 273, 823–829. doi:10.1098/rspb.2005.3385

Lee, K.P., Simpson, S.J., Clissold, F.J., Brooks, R., Ballard, J.W.O., Taylor, P.W., Soran, N., Raubenheimer, D., 2008. Lifespan and reproduction in *Drosophila*: new insights from nutritional geometry. Proc. Natl. Acad. Sci. U.S.A. 105, 2498–2503. doi:10.1073/pnas.0710787105

Li, J., Chen, W., Wu, J., Peng, W., An, J., Schmid-Hempel, P., Schmid-Hempel, R., 2012. Diversity of *Nosema* associated with bumblebees (*Bombus* spp.) from China. Int. J. Parasitol. 42, 49–61. doi:10.1016/j.ijpara.2011.10.005

Li, W., Chen, Y., Cook, S.C., 2018. Chronic *Nosema ceranae* infection inflicts comprehensive and persistent immunosuppression and accelerated lipid loss in host *Apis mellifera* honey bees. Int. J. Parasitol. 48, 433–444. doi:10.1016/j.ijpara.2017.11.004

Li, W., Evans, J.D., Li, J., Su, S., Hamilton, M., Chen, Y., 2017. Spore load and immune response of honey bees naturally infected by *Nosema ceranae*. Parasitol. Res. 116, 3265–3274. doi:10.1007/s00436-017-5630-8

Maklakov, A.A., Simpson, S.J., Zajitschek, F., Hall, M.D., Dessmann, J., Clissold, F., Raubenheimer, D., Bonduriansky, R., Brooks, R.C., 2008. Sex-Specific Fitness Effects of nutrient intake on reproduction and lifespan. Curr. Biol. 18, 1062–1066. doi:10.1016/j.cub.2008.06.059

Malysh, J.M., Ignatieva, A.N., Artokhin, K.S., Frolov, A.N., Tokarev, Y.S., 2018. Natural infection of the beet webworm *Loxostege sticticalis* L. (Lepidoptera: Crambidae) with three Microsporidia and host switching in *Nosema ceranae*. Parasitol. Res. doi:10.1007/s00436-018-5987-3

Martín-Hernández, R., Bartolomé, C., Chejanovsky, N., Le Conte, Y., Dalmon, A., Dussaubat, C., García-Palencia, P., Meana, A., Pinto, M.A., Soroker, V., Higes, M., 2018. *Nosema ceranae* in *Apis mellifera* : a 12 years postdetection perspective. Environ. Microbiol. 20, 1302–1329. doi:10.1111/1462-2920.14103

Martín-Hernández, R., Botías, C., Barrios, L., Martínez-Salvador, A., Meana, A., Mayack, C., Higes, M., 2011. Comparison of the energetic stress associated with experimental *Nosema ceranae* and *Nosema apis* infection of honeybees (*Apis mellifera*). Parasitol. Res. 109, 605–612. doi:10.1007/s00436-011-2292-9

Martín-Hernández, R., Higes, M., Sagastume, S., Juarranz, Á., Dias-Almeida, J., Budge, G.E., Meana, A., Boonham, N., 2017. Microsporidia infection impacts the host cell’s cycle and reduces host cell apoptosis. PLoS One 12, e0170183. doi:10.1371/journal.pone.0170183

Martín-Hernández, R., Meana, A., Prieto, L., Salvador, A.M., Garrido-Bailón, E., Higes, M., 2007. Outcome of colonization of *Apis mellifera* by *Nosema ceranae*. Appl. Environ. Microbiol. 73, 6331–8. doi:10.1128/AEM.00270-07

Mason, A.P., Smilanich, A.M., Singer, M.S., 2014. Reduced consumption of protein-rich foods follows immune challenge in a polyphagous caterpillar. J. Exp. Biol. 217, 2250–2260. doi:10.1242/jeb.093716

Mayack, C., Naug, D., 2009. Energetic stress in the honeybee *Apis mellifera* from *Nosema ceranae* infection. J. Invertebr. Pathol. 100, 185–8. doi:10.1016/j.jip.2008.12.001

Michener, C.D., 2000. The bees of the world, The Johns Hopkins University Press. Baltimore.

Mohr, K.I., Tebbe, C.C., 2006. Diversity and phylotype consistency of bacteria in the guts of three bee species (Apoidea) at an oilseed rape field. Environ. Microbiol. 8, 258–272. doi:10.1111/j.1462-2920.2005.00893.x

Perry, C.J., Søvik, E., Myerscough, M.R., Barron, A.B., 2015. Rapid behavioral maturation accelerates failure of stressed honey bee colonies. Proc. Natl. Acad. Sci. U. S. A. 112, 3427–32. doi:10.1073/pnas.1422089112

Piiroinen, S., Botías, C., Nicholls, E., Goulson, D., 2016. No effect of low-level chronic neonicotinoid exposure on bumblebee learning and fecundity. PeerJ 4, e1808. doi:10.7717/peerj.1808

Piiroinen, S., Goulson, D., 2016. Chronic neonicotinoid pesticide exposure and parasite stress differentially affects learning in honeybees and bumblebees. Proc. R. Soc. B Biol. Sci. 283, 20160246. doi:10.1098/rspb.2016.0246

Plischuk, S., Martín-Hernández, R., Prieto, L., Lucía, M., Botías, C., Meana, A., Abrahamovich, A.H., Lange, C., Higes, M., 2009. South American native bumblebees (Hymenoptera: Apidae) infected by *Nosema ceranae* (Microsporidia), an emerging pathogen of honeybees (*Apis mellifera*). Environ. Microbiol. Rep. 1, 131–5. doi:10.1111/j.1758-2229.2009.00018.x

Poissonnier, L.-A., Lihoreau, M., Gomez-Moracho, T., Dussutour, A., Buhl, J., 2018. A theoretical exploration of dietary collective medication in social insects. J. Insect Physiol. 106, 78–87. doi:10.1016/j.jinsphys.2017.08.005

Ponton, F., Wilson, K., Cotter, S.C., Raubenheimer, D., Simpson, S.J., 2011. Nutritional immunology: a multi-dimensional approach. PLoS Pathog. 7, 1–4. doi:10.1371/journal.ppat.1002223

Ponton, F., Wilson, K., Holmes, A., Raubenheimer, D., Robinson, K.L., Simpson, S.J., 2015. Macronutrients mediate the functional relationship between *Drosophila* and *Wolbachia*. Proc. R. Soc. B Biol. Sci. 282, 20142029–20142029. doi:10.1098/rspb.2014.2029

Porrini, M.P., Porrini, L.P., Garrido, P.M., de Melo e Silva Neto, C., Porrini, D.P., Muller, F., Nuñez, L.A., Alvarez, L., Iriarte, P.F., Eguaras, M.J., 2017. *Nosema ceranae* in south american native stingless bees and social wasp. Microb. Ecol. 2–5. doi:10.1007/s00248-017-0975-1

Porrini, M.P., Sarlo, E.G., Medici, S.K., Garrido, P.M., Porrini, D.P., Damiani, N., Eguaras, M.J., 2011. *Nosema ceranae* development in *Apis mellifera* : influence of diet and infective inoculum. J. Apic. Res. 50, 35–41. doi:10.3896/IBRA.1.50.1.04

Povey, S., Cotter, S.C., Simpson, S.J., Wilson, K., 2014. Dynamics of macronutrient self-medication and illness-induced anorexia in virally infected insects. J. Anim. Ecol. 83, 245–255. doi:10.1111/1365-2656.12127

Rapkin, J., Jensen, K., Archer, C.R., House, C.M., Sakaluk, S.K., Castillo, E. Del, Hunt, J., 2018. The geometry of nutrient space–based life-history trade-offs: sex-specific effects of macronutrient intake on the trade-off between encapsulation ability and reproductive effort in decorated crickets. Am. Nat. 191, 452–474. doi:10.1086/696147

Reddiex, A.J., Gosden, T.P., Bonduriansky, R., Chenoweth, S.F., 2013. Sex-Specific Fitness consequences of nutrient intake and the evolvability of diet preferences. Am. Nat. 182, 91–102. doi:10.1086/670649

Rinderer, T.E., Elliott, K.D., 1977. Worker honey bee response to infection with *Nosema apis*: Influence of diet. J. Econ. Entomol. 70, 431–433. doi:10.1093/jee/70.4.431ADDIN

Ruedenauer, F.A., Raubenheimer, D., Kessner-Beierlein, D., Grund-Mueller, N., Noack, L., Spaethe, J., Leonhardt, S.D. 2020. Best be(e) on low fat: linking nutrient perception, regulation and fitness. Ecol Lett. 23, 545–554. doi:10.1111/ele.13454

Roberts, K.E., Hughes, W.O.H., 2015. Horizontal transmission of a parasite is influenced by infected host phenotype and density. Parasitol. 142, 395–405. doi:10.1017/S0031182014001243

Schmid-Hempel, P., 1998. Parasites in social insects. Princeton University Press, Princeton, New Jersey.

Schmid-Hempel, R., Tognazzo, M., 2010. Molecular divergence defines two distinct lineages of *Crithidia bombi* (Trypanosomatidae), parasites of bumblebees. J. Eukaryot. Microbiol. 57, 337–45. doi:10.1111/j.1550-7408.2010.00480.x

Semaniuk, U., Feden’ko, K., Yurkevych, I.S., Storey, K.B., Simpson, S.J., Lushchak, O., 2018. Within-diet variation in rates of macronutrient consumption and reproduction does not accompany changes in lifespan in *Drosophila melanogaster*. Entomol. Exp. Appl. 166, 74–80. doi:10.1111/eea.12643

Therneau, T.M., Grambsch, P.M., 2000. Modeling survival data: extending the cox model, statistics for biology and health. New York: Springer

Tritschler, M., Vollmann, J.J., Yañez, O., Chejanovsky, N., Crailsheim, K., Neumann, P., 2017. Protein nutrition governs within host race of honey bee pathogens. Sci. Rep. 7, 14988. doi:10.1038/s41598-017-15358-w

Vaudo, A.D., Patch, H.M., Mortensen, D.A., Tooker, J.-F., Grozinger, C.M., 2016. Macronutrient ratios in pollen shape bumble bee (*Bombus impatiens*) foraging strategies and floral preferences. Proc Natl Acad Sci USA. 113, E4035–E4042. doi:10.1073/pnas.1606101113

Van der Zee, R., Gómez-Moracho, T., Pisa, L., Sagastume, S., García-Palencia, P., Maside, X., Bartolomé, C., Martín-Hernández, R., Higes, M., 2014. Virulence and polar tube protein genetic diversity of *Nosema ceranae* (Microsporidia) field isolates from Northern and Southern Europe in honeybees (*Apis mellifera iberiensis*). Environ. Microbiol. Rep. 6, 401–413. doi:10.1111/1758-2229.12133

Venables, W.N., Ripley, B.D., 2002. Modern applied statistics with S. New York: Springer

Wittner, M., Weiss, L., 1999. The Microsporidia and microsporidiosis. American Society of Microbiology. doi:10.1128/9781555818227

Wolf, S., McMahon, D.P., Lim, K.S., Pull, C.D., Clark, S.J., Paxton, R.J., Osborne, J.L., 2014. So near and yet so far: harmonic radar reveals reduced homing ability of *Nosema* infected honeybees. PLoS One 9, e103989. doi:10.1371/journal.pone.0103989

Zheng, H.Q., Lin, Z.G., Huang, S.K., Sohr, A., Wu, L., Chen, Y.P., 2014. Spore loads may not be used alone as a direct indicator of the severity of *Nosema ceranae* infection in honey bees *Apis mellifera* (Hymenoptera:Apidae). J. Econ. Entomol. 107, 2037–2044. doi:10.1603/ec13520

