## Supplementary Materials for "Artificial diets modulate infection rates by *Nosema ceranae* in bumblebees" (from another version of this preprint: version 1, 2020-02-27)

**Table 1. Number of bumblebees positive to *N. ceranae* in a PCR and showing spores at different days post exposure (dpe) per spore dosages and diet treatment**. LP: low-protein diet; HP: high-protein diet.

| | | 150K | | | 300K | | |
|---|---|---|---|---|---|---|---|
| **Diet** | **Dpe** | **Dead** | **PCR positive** | **Spores** | **Dead** | **PCR positive** | **Spores** |
| LP | 1 | 0 | 0 | 0 | 0 | 0 | 0 |
| LP | 2 | 3 | 3 | 0 | 2 | 2 | 0 |
| LP | 3 | 5 | 2 | 1 | 3 | 2 | 1 |
| LP | 4 | 5 | 3 | 1 | 7 | 6 | 1 |
| LP | 5 | 4 | 0 | 0 | 7 | 1 | 0 |
| LP | 6 | 6 | 3 | 1 | 4 | 2 | 2 |
| LP | 7 | 2 | 1 | 0 | 3 | 2 | 1 |
| LP | 8 | 3 | 1 | 0 | 4 | 2 | 0 |
| LP | 9 | 3 | 1 | 0 | 0 | 0 | 0 |
| LP | 10 | 0 | 0 | 0 | 2 | 2 | 0 |
| LP | 11 | 1 | 1 | 0 | 0 | 0 | 0 |
| LP | 12 | 0 | 0 | 0 | 0 | 0 | 0 |
| LP | 13 | 0 | 0 | 0 | 0 | 0 | 0 |
| LP | 14 | 1 | 0 | 0 | 0 | 0 | 0 |
| LP | 15 | 1 | 1 | 0 | 1 | 1 | 0 |
| LP | 16 | 0 | 0 | 0 | 1 | 1 | 0 |
| LP | 17 | 0 | 0 | 0 | 1 | 0 | 0 |
| LP | 18 | 0 | 0 | 0 | 1 | 1 | 0 |
| LP | 19 | 2 | 1 | 0 | 0 | 0 | 0 |
| LP | 20 | 6 | 6 | 4 | 9 | 9 | 4 |
| LP | **TOTAL** | ***42*** | ***23*** | ***7*** | ***45*** | ***31*** | ***9*** |
| HP | 1 | 0 | 0 | 0 | 0 | 0 | 0 |
| HP | 2 | 2 | 1 | 1 | 2 | 1 | 0 |
| HP | 3 | 4 | 3 | 0 | 3 | 3 | 3 |
| HP | 4 | 2 | 1 | 0 | 4 | 2 | 0 |
| HP | 5 | 4 | 0 | 0 | 5 | 4 | 1 |
| HP | 6 | 5 | 2 | 2 | 3 | 1 | 0 |
| HP | 7 | 5 | 0 | 0 | 3 | 1 | 1 |
| HP | 8 | 13 | 6 | 1 | 9 | 1 | 0 |
| HP | 9 | 4 | 0 | 0 | 3 | 0 | 1 |
| HP | 10 | 3 | 0 | 0 | 6 | 2 | 0 |

| | | | | | | | |
|---|---|---|---|---|---|---|---|
| HP | 11 | 2 | 0 | 0 | 1 | 0 | 0 |
| HP | 12 | 3 | 1 | 0 | 0 | 0 | 0 |
| HP | 13 | 0 | 0 | 0 | 0 | 0 | 0 |
| HP | 14 | 0 | 0 | 0 | 0 | 0 | 0 |
| HP | 15 | 0 | 0 | 0 | 1 | 0 | 0 |
| HP | 16 | 0 | 0 | 0 | 0 | 0 | 0 |
| HP | 17 | 0 | 0 | 0 | 0 | 0 | 0 |
| HP | 18 | 0 | 0 | 0 | 0 | 0 | 0 |
| HP | 19 | 0 | 0 | 0 | 0 | 0 | 0 |
| HP | 20 | 0 | 0 | 0 | 1 | 0 | 0 |
| HP | **TOTAL** | ***47*** | ***14*** | ***4*** | ***41*** | ***15*** | ***6*** |

**Table 2. Tukey post-hoc test to account for pairwise comparisons between bumblebees in different diets and spore treatments**. Groups are defined by diet.dosage.infection. Diet: C (low-protein), P (high-protein). Dosage: 0, 150, 300. Infection: 0 (not-infected), 1 (infected).

| Group 1 | | Group 2 | Estimate | SE | z | p | |
|---|---|---|---|---|---|---|---|
| C.150.0 | - | C.0.0 | -110.587 | 0.14964 | -7.390 | <0.01 | *** |
| C.150.0 | - | P.0.0 | -0.07837 | 0.12572 | -0.623 | 1.000 | |
| C.150.1 | - | C.0.0 | -0.49910 | 0.15674 | -3.184 | 0.0554 | . |
| C.150.1 | - | P.0.0 | 0.52840 | 0.13485 | 3.918 | <0.01 | ** |
| C.150.1 | - | C.150.0 | 0.60677 | 0.15356 | 3.951 | <0.01 | ** |
| C.150.1 | - | P.150.0 | 0.59391 | 0.14048 | 4.228 | <0.01 | ** |
| C.150.1 | - | C.300.0 | 0.69054 | 0.16983 | 4.066 | <0.01 | ** |
| C.150.1 | - | P.300.0 | 0.44123 | 0.14369 | 3.071 | 0.0774 | . |
| C.300.0 | - | C.0.0 | -118.964 | 0.16624 | -7.156 | <0.01 | *** |
| C.300.0 | - | P.0.0 | -0.16214 | 0.14513 | -1.117 | 0.9927 | |
| C.300.0 | - | C.150.0 | -0.08377 | 0.16228 | -0.516 | 1.000 | |
| C.300.0 | - | P.150.0 | -0.09663 | 0.15034 | -0.643 | 1.000 | |
| C.300.1 | - | C.0.0 | -0.44593 | 0.14664 | -3.041 | 0.0845 | . |
| C.300.1 | - | P.0.0 | 0.58157 | 0.12332 | 4.716 | <0.01 | *** |
| C.300.1 | - | C.150.0 | 0.65995 | 0.14374 | 4.591 | <0.01 | *** |
| C.300.1 | - | P.150.0 | 0.64709 | 0.12947 | 4.998 | <0.01 | *** |
| C.300.1 | - | C.300.0 | 0.74372 | 0.16100 | 4.619 | <0.01 | *** |
| C.300.1 | - | P.300.0 | 0.49441 | 0.13298 | 3.718 | <0.01 | ** |
| C.300.1 | - | C.150.1 | 0.05318 | 0.15190 | 0.350 | 1.000 | |
| C.300.1 | - | P.150.1 | 0.79892 | 0.16081 | 4.968 | <0.01 | *** |
| P.0.0 | - | C.0.0 | -102.750 | 0.12842 | -8.001 | <0.01 | *** |
| P.150.0 | - | C.0.0 | -109.301 | 0.13448 | -8.127 | <0.01 | *** |
| P.150.0 | - | P.0.0 | -0.06551 | 0.10885 | -0.602 | 1.000 | |
| P.150.0 | - | C.150.0 | 0.01286 | 0.13169 | 0.098 | 1.000 | |
| P.150.1 | - | C.0.0 | -124.485 | 0.16532 | -7.530 | <0.01 | *** |
| P.150.1 | - | P.0.0 | -0.21735 | 0.14480 | -1.501 | 0.9308 | |
| P.150.1 | - | C.150.0 | -0.13897 | 0.16241 | -0.856 | 0.9993 | |
| P.150.1 | - | P.150.0 | -0.15183 | 0.15006 | -1.012 | 0.9969 | |
| P.150.1 | - | C.300.0 | -0.05520 | 0.17787 | -0.310 | 1.000 | |
| P.150.1 | - | P.300.0 | -0.30451 | 0.15307 | -1.989 | 0.6703 | |
| P.150.1 | - | C.150.1 | -0.74574 | 0.16975 | -4.393 | <0.01 | *** |
| P.300.0 | - | C.0.0 | -0.94034 | 0.13815 | -6.807 | <0.01 | *** |
| P.300.0 | - | P.0.0 | 0.08716 | 0.11305 | 0.771 | 0.9998 | |
| P.300.0 | - | C.150.0 | 0.16554 | 0.13502 | 1.226 | 0.9843 | |
| P.300.0 | - | P.150.0 | 0.15268 | 0.11973 | 1.275 | 0.9787 | |

| | | | | | | | |
|---|---|---|---|---|---|---|---|
| P.300.0 | - | C.300.0 | 0.24931 | 0.15327 | 1.627 | 0.8847 | |
| P.300.1 | - | C.0.0 | -135.445 | 0.15933 | -8.501 | <0.01 | *** |
| P.300.1 | - | P.0.0 | -0.32695 | 0.13784 | -2.372 | 0.3920 | |
